# An 11-bit High Dynamic Range (HDR) Luminance Display and Its Use to Discover Contextual Mechanisms of Real-World Luminance Normalization for Visual Acuity and Target Discrimination

**DOI:** 10.1101/718437

**Authors:** Chou P Hung, Chloe Callahan-Flintoft, Paul D Fedele, Kim F Fluitt, Onyekachi Odoemene, Barry D Vaughan, Anthony J Walker, Matthew M Jaswa, Min Wei, Andre V Harrison

**Affiliations:** Human Research and Engineering Directorate, CCDC Army Research Laboratory; DCS Corporation; Computational and Information Sciences Directorate, CCDC Army Research

**Keywords:** human-autonomy teaming, human-autonomy integration, human variability, real-world behavior, translational neuroscience, automatic target recognition (ATR), resilience, visual search

## Abstract

Luminance can vary widely when scanning across a scene, by up to 10^9 to 1, requiring multiple normalizing mechanisms spanning from the retina to cortex to support visual acuity and recognition. Vision models based on standard dynamic range luminance contrast ratios below 100 to 1 have limited ability to generalize to real-world scenes with contrast ratios over 10,000 to 1 (high dynamic range [HDR]). Understanding and modeling brain mechanisms of HDR luminance normalization is thus important for military applications, including automatic target recognition, display tone mapping, and camouflage. Yet, computer display of HDR stimuli was until recently unavailable or impractical for research. Here we describe procedures for setup, calibration, and precision check of an HDR display system with over 100,000 to 1 luminance dynamic range (650–0.0065 cd/m^2), pseudo 11-bit grayscale precision, and 3-ms temporal precision in the MATLAB/Psychtoolbox software environment. The setup is synchronized with electroencephalography and IR eye-tracking measurements. We report measures of HDR visual acuity and the discovery of a novel phenomenon—that abrupt darkening (from 400 to 4 cd/m^2) engages contextual facilitation, distorting the perceived orientation of a high-contrast central target. Surprisingly, the facilitation effect depended on luminance similarity, contradicting both classic divisive and subtractive models of contextual normalization.

## 1. Introduction

### 1.1 The Challenge of High Dynamic Range (HDR) Luminance for Human and Machine Vision

Resilient and intuitive machine vision is a critical capability for autonomous teaming and is therefore key to multiple Army modernization priorities, yet the real-world complexity of this challenge is persistently underestimated because vision feels so effortless. Vision is an inherently ambiguous process of estimating and predicting the true 3-D world from a 2-D retinal image, and the apparent ease of vision is belied by the fact that nearly half of the brain is devoted to visual processing. Depending on the context of a visual scene, almost any luminance can appear as any shade of gray. This is because the luminance of the brightest and darkest areas can vary by a factor of up to 1 billion to 1 (Fig. 1), whereas the surface reflectance information that is useful for estimating object shape and identity typically varies by a factor of only 20 to 1 (i.e., 4% to 80% of the light illuminating the surface (Gilchrist et al. 1999). Even slight (<0.5%) changes in illumination, as due to variations in atmospheric haze or sun position, can produce large (50%) average changes in luminance in a natural scene; for example, due to cast shadows, foliage, and anisotropic reflectance effects such as microshadows (Moore 2010; Foster and Amano 2019). Large dynamic changes, such as from flare effects and wind blowing on the leaves in a forest canopy (Marathe et al. 2017), can disrupt navigation and targeting algorithms that are over-reliant on texture patterns and on static luminance and illumination to compute optic flow. For autonomous ground vehicles, navigation is problematic because luminance normalization is not yet solved for recognizing distant targets (e.g., potholes, buried explosives) under multiple layers of optic flow, especially at high speed and under degraded and novel conditions. These pose serious longstanding challenges to the real-world credibility and acceptability of autonomous maneuver and targeting, with potentially catastrophic consequences, for example, from false positive and false negative misclassifications.

**Fig. 1.**
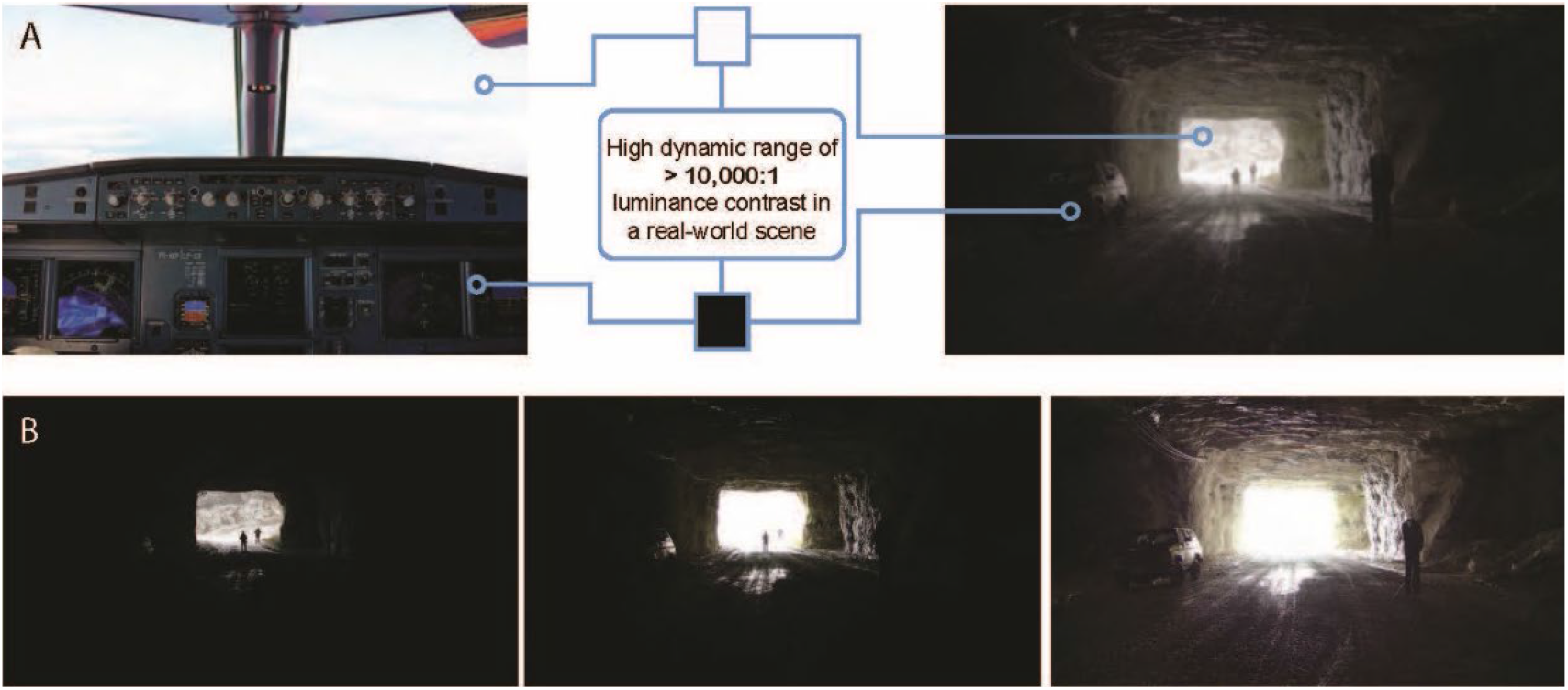
A) Two examples of HDR luminance in naturalistic scenes: cockpit view (left) and view of a cave opening (right). They are examples of commonly encountered environments where combinations of indoor and outdoor luminance can exceed 10,000-to-1 max-to-min luminance ratio. The scene at top right is a blended image across multiple exposures, illustrating our ability to see multiple targets (3 uniforms and 1 car) across vast luminance differences in the same view. Actual views of these scenes appear even more vivid because of the brain’s luminance normalization processes. B) Standard dynamic range 8-bit cameras are highly dependent on exposure, and machine vision systems that are highly dependent on texture patterns can easily miss targets due to under- or overexposure. Exposure examples are at 10× increments.

For efficient and resilient recognition behavior, the human visual system has many automatic mechanisms to estimate reflectance and 3-D shape, as demonstrated by myriad examples of brightness illusions (Adelson 2000; Motoyoshi et al. 2007; Bach 2019). Our understanding of these mechanisms, as quantified by models of normalization and visual salience, is limited because it is based on studies using displays with standard dynamic range (SDR) luminance (100-to-1 luminance ratio between the brightest and darkest pixels). Recent reports show that key theories, such as Wallach’s ratio rule that the apparent lightness of a surface depends on the ratio of its luminance to the background, break down at HDR luminance (over 10,000-to-1 luminance ratio). To investigate and understand these mechanisms, and thereby improve the real-world performance of vision models, it is necessary to develop research platforms that include HDR displays.

To achieve illumination resilient recognition, biological visual systems gradually transform the retinal image from a luminance-based representation to reflectance-approximated feature-based representations (Zhou et al. 2000; Janssen et al 2001; Wachtler et al. 2003; Roe et al. 2012). Evidence from anatomy, electrophysiology, and behavior show that this is supported by a hierarchy of over 30 visual cortical areas that combine recurrent and lateral processes with feedback mechanisms for contextual normalization (Ramsden et al. 2001; Tanigawa et al. 2005). The brain areas that support normalization are highly specialized, including specific domains for luminance and color processing within multiple visual cortical areas including the primary visual cortex (Area V1, the first brain area to receive input from the retina and thalamus) (Schroeder et al. 1998; Conway et al. 2007; Wang et al. 2007; Lim et al. 2009; Kremkow et al. 2016). The circuitry underlying these processes is less understood, but there is evidence for tight coupling between processes for luminance/color and processes for form perception, including feedback circuitry from the last stage of the visual “form” pathway to V1 (Ts’o et al. 2001; Clavagnier et al. 2004; Hung et al. 2007, 2010). Rather than simply remapping the color histogram or normalizing an image for nearby luminance, automatic mechanisms are thought to depend on factors such as co-linearity, co-planarity, junctions, feature grouping, and transparency issues such as smoke and rain (Zucker et al. 1988; Anderson 1997; Adelson 2000). Biologically driven models are increasingly capable of explaining visual illusions (Blakeslee and McCourt 2004; Li 2011) and predicting gaze patterns based on saliency (Borji 2018; Kummerer et al. 2017), but they are data-limited to SDR images and require HDR experimentation to extend their generalizability to real-world vision.

### 1.2 HDR Research to Understand Vision under Real-World Illumination

Historically, the unavailability of HDR displays has been a primary impediment to understanding how the brain processes real-world luminance, underlying a major gap between biological and machine vision. Early efforts to understand HDR vision were based on hand-constructed scenes (Zdravković et al. 2012; Brémond et al. 2010), and it was only recently that computerized displays became available for more-sophisticated studies. A few studies have used custom-developed stacked liquid crystal displays (LCDs) for HDR, but such systems are difficult to replicate and lack sufficient spatial resolution and uniformity for our studies. Commercial and research displays are typically limited to less than 1000-to-1 luminance contrast ratio and 8- to 10-bit depth (i.e., 256 to 1024 shades of gray).

Here we describe the setup, calibration, and precision testing of a system with more than 100,000-to-1 luminance contrast ratio and pseudo 11-bit depth (2048 shades of gray), enabling the study of the visual system’s HDR luminance normalization processes. We also describe two studies we conducted with this system to understand how luminance dynamics affect visual letter acuity and how luminance and surround context together affect target discrimination. For the letter acuity study (Fig. 2A), our system allowed us to present smaller letters by avoiding potential spatial misalignment and inhomogeneity of stacked LCD displays. We extended a previously developed computerized Logarithm of the minimum-angle-of-resolution (LogMAR) letter acuity test with (standard) static luminance (Stewart et al. 2006) by adding HDR luminance dynamics. Specifically, we tested the effect of bright (25× to 100× luminance) flashes on the acuity of dark letters at different background and letter luminance contrasts, as might be encountered when switching gaze between outside viewing and reading an instrument panel. In addition to being an easily understood measure tied to standard LogMAR acuity testing, this approach could also be useful to investigate medical phenomena such as age-related decline in pupil accommodation.

**Fig. 2.**
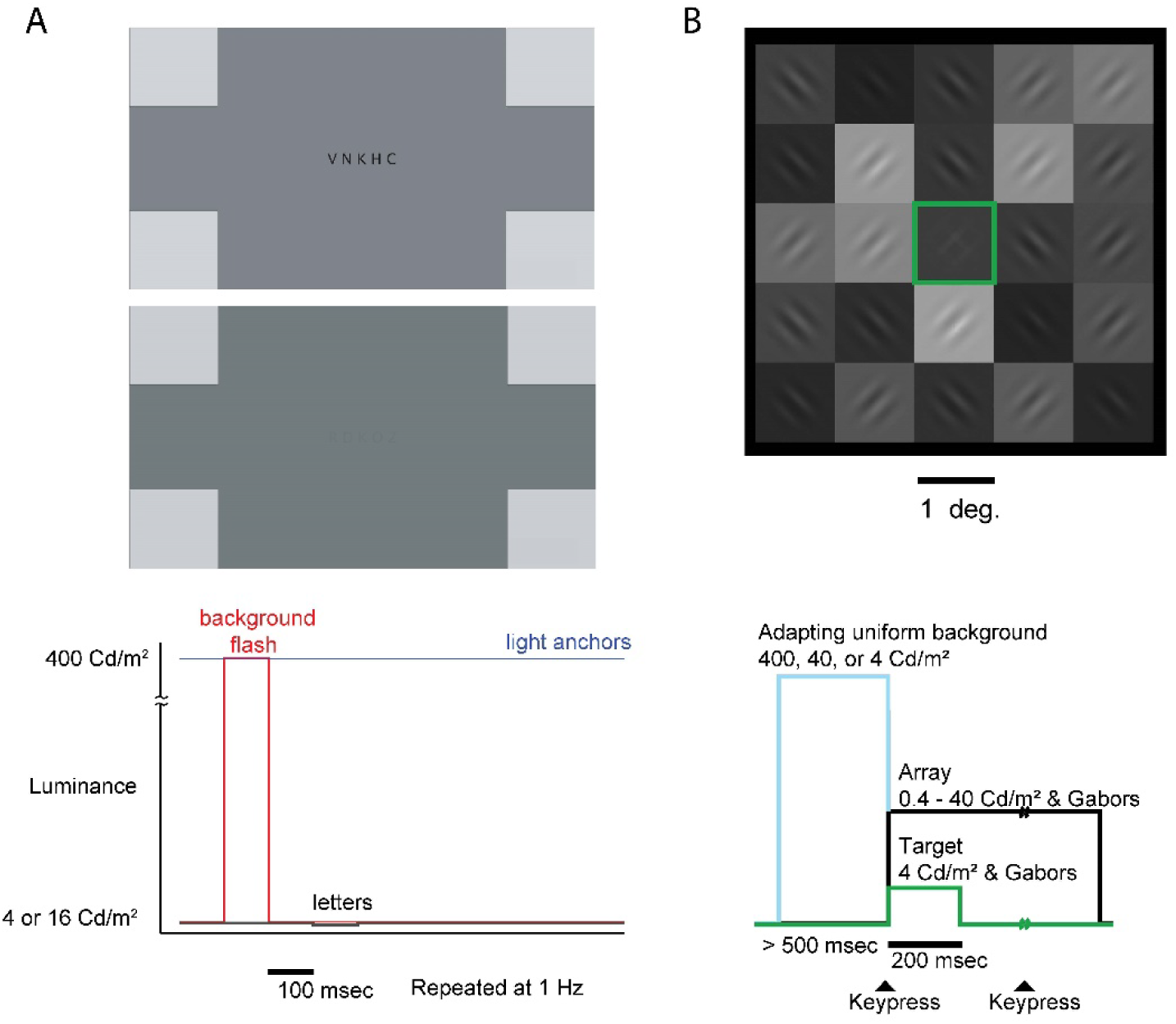
A) Dynamic Luminance Visual Acuity Task (Task 1), a variant of the classic LogMAR chart. We manipulated background luminance and letter contrasts and added a background flash preceding letter onset; (top left) two examples of the letter arrays presented to participants, shown at letter Weber contrast levels of 0.999 and 0.25; (bottom left) timecourse of the flash and letter presentations. B) Schematic of the HDR Luminance Target Discrimination Task (Task 2), a two-alternative forced choice task in which the subject reported the orientation of the central target (highlighted in green) as one of two Gabors having higher contrast; (top right) example of the 5 × 5 Gabor array; (bottom right) timecourse of adapting uniform background, array, and target presentations and keypresses.

For the HDR target discrimination study (Fig. 2B), our aim was to investigate contextual mechanisms for luminance normalization by testing for interactions between orientation and HDR luminance processing. Based on recent reports on HDR luminance normalization (Radonjic et al. 2011; Allred et al. 2012) and on single neuron receptive field properties in V1, we speculated that at least a 10,000-to-1 contrast ratio and better than 0.06° pixel resolution (see Methods, Section 2) would be required to observe HDR-specific interactions between luminance and orientation processing in our task. This requirement is beyond the capabilities of SDR displays but is well within the 100,000-to-1 contrast ratio and 0.02° pixel resolution capability of our HDR display system.

Previous reports of contextual orientation effects found that flankers drive a facilitating response (making a co-oriented target easier to detect) if the targets were low contrast, and this was initially attributed to horizontal fibers linking V1 neurons that prefer the same orientation (Ts’o et al. 1986). However, at higher contrast a co-oriented target becomes more difficult to detect than an orthogonal target, an effect that is consistent with suppression of the target visibility or assimilation of the target to surrounding co-oriented patterns, possibly due to feedback from higher pattern-sensitive cortical areas. Both of these phenomena have also been attributed to the balance of local recurrent excitatory and inhibitory mechanisms in V1, but they have thus far only been investigated for static luminance displays and uniform patch luminance (Polat and Sagi 1993, 2006; Li 1998, 2011; Polat et al. 1998; Chen and Tyler 2001, 2002; Chen et al. 2001).

We reasoned that in naturalistic vision, our gazes often shift across regions with large (100×) differences in luminance. How does the visual system normalize quickly, perhaps even predictively (peri-saccade), across large luminance changes, and can we discover such a normalizing mechanism, for example, linking luminance and form vision, by observing how shape perception is altered when shifting one’s gaze from light to dark areas of the visual scene during visual search? Would strong darkening lead to contextual facilitation, even for high-contrast targets, or suppression/assimilation? Also, would the contextual effect be driven by the brightest flankers, consistent with models of recurrent excitation/inhibition and divisive or subtractive normalization, or would the effects be driven by the flankers that are most similar in luminance to the target, consistent with assimilation or feedback from higher brain areas? To answer these questions, we took the novel approach of combining 1) a luminance transition of 1×, 10×, or 100×, 2) the 5 × 5 array pattern of recent HDR luminance studies (Radonjic et al. 2011), and 3) oriented lines (“flankers”) that surround the target and affect target visibility (classic contextual orientation effects discovered with static SDR displays (Chen and Tyler 2002), and we measured how these contextual HDR luminance manipulations affect orientation discrimination of a central target.

## 2. Methods

### 2.1 Subjects

Nine subjects (six male) 18 to 70 years old with normal or corrected-to-normal binocular color vision participated in all experiments. Potential subjects were excluded if they self-reported that they, their parents, or their siblings had photosensitive epilepsy, or that they previously had head trauma or other disorders thought to be associated with excitatory/inhibitory balance (epilepsy, schizophrenia, autism, depression, attention deficit hyperactivity disorder, etc.) (Jiang et al. 2013), atypical brain development, or used mind-altering drugs in the past week. Potential subjects were also screened via the Canadian Longitudinal Study on Aging – Epilepsy Algorithm (Keezer et al. 2014). All experiments were conducted in the MIND lab at the CCDC Army Research Laboratory at Aberdeen Proving Ground, Maryland, according to a protocol approved by the Army’s Human Research Protection Program.

### 2.2 Vision Screening

Prior to beginning experimental tasks, subjects were screened for normal or corrected-to-normal (at least 20/40) visual acuity and normal color vision via a Titmus i500 Vision Screener (Lipin-Dietz 2012).

### 2.3 HDR Display and Eye Tracking

All images were projected from a JVC DLA-RS600U 4K Reference Projector (software version u83.2, PS version 100310) and displayed biocularly on an HD projection screen (Fig. 3; Da-Lite 37700V Cinema Contour screen with HD surface, “HD Progressive 1”, gain = 1.10 at 0° and 1.00 at 20°, half angle = 85°, gloss at 75° = 24, color shift at 60° = 3%). To maximize the contrast ratio, the projector was set to the smallest zoom and the following settings: Color Profile “Reference”, Color Temp “6500K”, Gamma “D”, MPC Level 4K e-shift “ON”, Original Resolution “Auto”, Enhance “0”, Dynamic contrast “0”, Smoothing “0”, NR “0”, Blur Reduction Clear Motion Drive “High”, Motion Enhance “Off”’, Contrast “0”, Brightness “0”, Color “20”, Tint “−5”, Input “HDMI-1”, Source “1080p 60”, Deep Color “12 bit”, and Color Space “RGB” (red-green-blue). Although the projector, graphics card, and HDMI cable supported 12-bits-per-channel bit depth, the overall system was software-limited to 10.7-bit depth. Images spanned 1920 × 1080 pixels in resolution (48.7 × 27.3 cm, w × h) and were observed from a chinrest-stabilized viewing distance of 78 cm, thus spanning 38.6° × 20.5° viewing angle with pixel size 0.020° × 0.019°. Gaze and pupil size were tracked monocularly via an infrared eye tracker (EyeLink 1000 Plus), synchronized via Lab Streaming Layer software (Swartz Center for Computational Neuroscience, UCSD) (Kothe 2014). To maintain a constant peak luminance in the visual field, all tasks included static 400- cd/m^2^ “light anchors” (Gilchrist et al. 1999) sized 1° × 1° at the four corners of the screen.

**Fig. 3.**
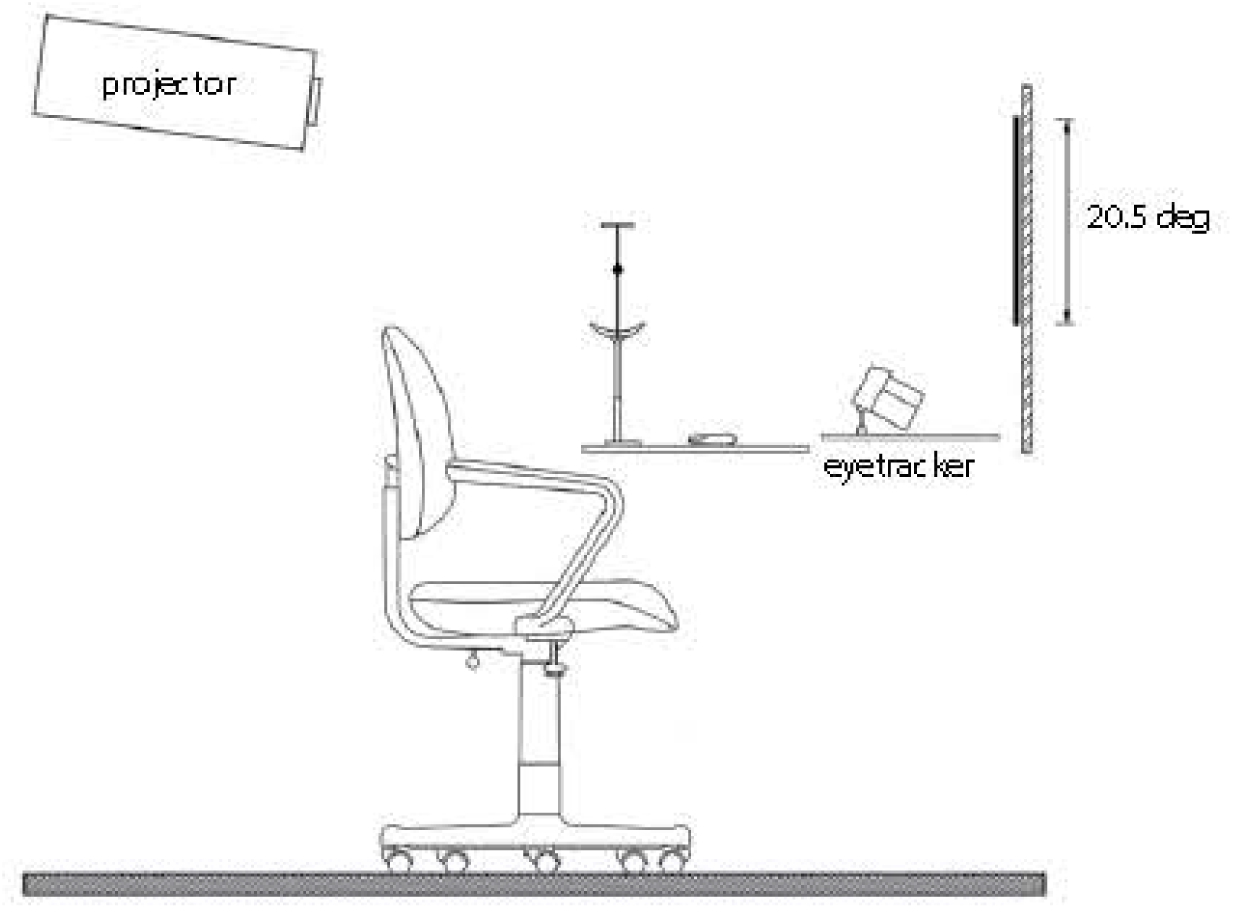
Layout of projector, screen, eye tracker, and chinrest

Images were displayed at 60 Hz and pseudo 11 bits (10.7 bits, i.e., 11 bits red, 11 bits green, but only 10 bits blue, because all the color information needed to fit into 32 bits) precision via a framebuffer procedure using Psychtoolbox 3.0 (Kleiner et al. 2007) for GNU/Linux X11 software (version 3.0.14; build date: 2017 May 8), running under MATLAB 64-bit version 2016b on Ubuntu 16.04 (seen by Psychtoolbox as Linux version 4.4.0-31-generic). Because 11-bit precision in Psychtoolbox 3.0 requires graphics cards supporting the AMD Hawaii PRO GL (DRM 2.43.0 / 4.4.0-31-generic, LLVM 3.8.0) (Kleiner 2017) (i.e., the Radeon/Fire cards of the “Sea Islands” family [after 2014]), we used the AMD FirePro W8100 graphics card. We validated a second system running OSX and a FirePro W5100 graphics card. We also tested other graphics cards outside the “Sea Islands” family (VisionTek RADEON R9 280X, MSI Twin Frozr III, and Nvidia Quadro) and found that, as expected, they were limited to 8 or 10 bits per channel (bpc). We used the Psychtoolbox command *PsychImaging(‘AddTask’,‘General’,‘EnableNative11BitFramebuffer’)* to disable and bypass the hardware’s Gamma color lookup table (“clut”) and switch the framebuffer into 11-bpc mode. We applied an 11-bit grayscale Gamma correction by measuring the luminance via a spectrophotometer (Photo Research PR-745) at over 75 luminance indices, averaging across two to five repeated measurements per index, and applying log-linear interpolation. The resulting linearized Gamma spanned a range of 636.4 (u,v = 0.1953, 0.3199; x,y = 0.3200, 0.3494, 6037K) to 0.006055 cd/m^2^, for a maximum contrast ratio of more than 100,000 to 1 in a single image (static projector iris; we did not assess the manufacturer’s stated capability of 1.5M contrast ratio with dynamic iris).

### 2.4 Projector Calibration

We calibrated the projector by measuring the luminance of a uniform full screen at over 75 luminance color indices (setting the RGB guns to equal values), averaging across multiple measurements per color index. The photometer sampled from the projector screen, rather than directly from the projector output, and this difference might explain why we did not observe the projector manufacturer’s specified maximum contrast ratio of 150,000 to 1. To reduce the noise effects of drift, we allowed the projector to warm up for 30 min before calibration, and we repeatedly measured at each color index before moving to the next color index. We used different photometer settings depending on the light level, measuring smaller 0.5° diameter and 2-nm-wide samples for the brightest indices and larger 2° diameter and 4-nm-wide samples (and ~15 s averaging per sample) for the darkest indices. To ensure that the calibration was correct for the entire range, we measured more samples at the steepest parts of the curve (e.g., 0.01 to 0.05 cd/m^2^) and near inflection points. We log-linearly interpolated the luminance for indices that were not measured, and the resulting Gamma curve is plotted in Fig. 4A. We observed a rightward shift of approximately150 color indices in this Gamma curve on the OSX platform (Mac Pro 3.2, model A1289, server mid-2010 family), which resulted in more dark pixels for the lowest color indices and a smaller range of ceiling effect, but otherwise no change to the Gamma curve. All experiments were based on the Ubuntu platform. The curve shows a steep rise in luminance for the lowest indices and a ceiling effect, resulting in an effective range of luminances from 0.006055 to 636.4 cd/m^2^ (uncorrected color indices from 16 to 1728). This range spans mesopic vision (0.001 to 3 cd/m^2^, when both cones and rods are required to support vision) to the lower end of photopic vision (10 to 10^8^ cd/m^2^), consistent with seeing in mixed indoor/outdoor environments and in twilight (e.g., nighttime street and outdoor lighting and aviation lighting).

**Fig. 4.**
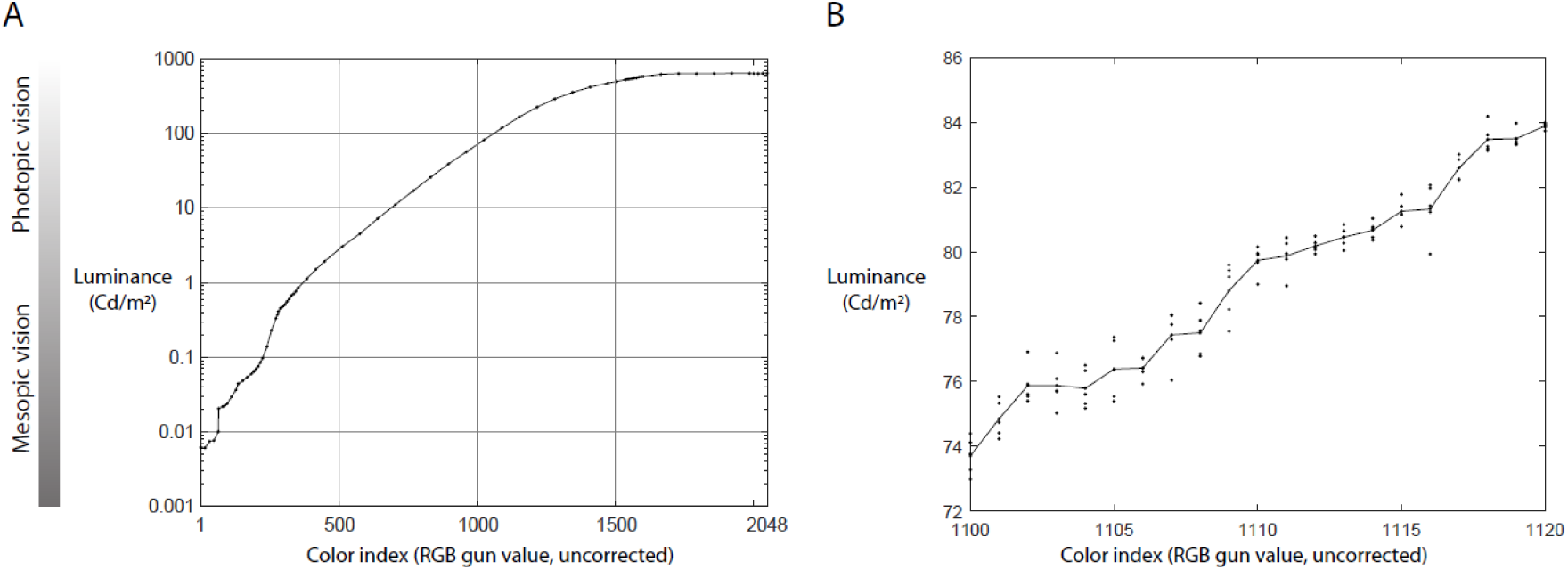
Luminance calibration and 11-bit precision check. A) Luminance calibration on the Ubuntu system with an AMD FirePro W8100 graphics card. B) Precision check on an OSX system and an AMD FirePro W5100 graphics card. Calibration curves were horizontally shifted but otherwise identical across the two systems.

### 2.5 Projection Luminance Precision

To our knowledge, we are the first to validate the 10.7-bits-per-channel precision of Psychtoolbox 3 via careful photometer measurements, due to lack of suitable display hardware at developer sites. We optimized for both display precision and photometer sensitivity by testing a range of uncorrected color indices (uniform RGB gun values) ranging from 1100 to 1120, at nearly three-quarters of the maximum luminance capability and the smoothest part of the Gamma curve. We measured five repeated samples per color index, completing measurements at each color index before moving on to the next to avoid slow drift effects on repeated measurements. The averaged measurements generally increased steadily, consistent with 10.7-bit precision, without obvious stairstepping that would indicate 10-bit precision or lower.

### 2.6 Display Timing Delay

To measure the timing offsets and precision between the MATLAB task control timestamps and when the images were actually updated on the projection screen, we ran a test block (398 trials) with a photodiode pointed toward the projector, positioned in the background part of the image just below the 5 × 5 array. The output of the photodiode was sent to Lab Streaming Layer via a custom Arduino device. Comparing the timestamp of when the code was executed to display the blank screen, to the time at which the photodiode registered the increase in luminance, showed an average projector display lag of 137 ± 2.8 ms. A similar comparison, this time for the blank’s offset (background darkening immediately followed by array and target onsets) showed an average lag of 162 ± 4.9 ms. We speculate that this difference of 24.4 ± 1.5 ms, just over one video frame at 60-Hz frame rate, is related to two additional MATLAB image updates for the array and target sent at blank offset. Both of these average lags were incorporated when forming epochs in the electroencephalography (EEG) data.

### 2.7 Experiment 1: Dynamic Luminance Visual Acuity Task

Visual acuity was measured via a modified Early Treatment Diabetic Retinopathy Study (ETDRS) Logarithm of the minimum-angle-of-resolution LogMAR chart presented via MATLAB, based on code modified from Stewart et al. 2006. An example of the letter display is shown in Fig. 2, Task 1. For each trial, a set of five letters was presented, pseudorandomly chosen from a predetermined pool of five-letter sets that were balanced for difficulty. The set of letters was repeatedly presented for 0.1 s ON and 0.9 s OFF until the subject read the letters aloud and the experimenter recorded the number of correct letters (0 to 5). A block comprised 20 trials, spanning letter sizes 3.125 to 50 minutes of arc (arcmin). On one block, dark letters (0.006 cd/m^2^) were presented against a background of 16 cd/m^2^. On four blocks, dark letters were presented against a uniform background of 4 or 16 cd/m^2^, at Weber contrasts of 0.43 and 0.25 for each background (e.g., for the 16 cd/m^2^ background, the letters were 9.2 cd/m^2^ for 0.43 contrast and 12.1 cd/m^2^ for 0.25 contrast). On five additional blocks, each presentation was preceded by a 400-cd/m^2^ flash of duration 0.1 s (i.e., 25× or 100× the background luminance). The flashed blocks were presented first in order of decreasing background luminance and contrast followed by the nonflashed blocks in the same sequence.

### 2.8 Experiment 2: HDR Luminance Target Discrimination Task

Task 2 was a two-alternative forced-choice target discrimination task comprising five experimental blocks. Across all blocks, the target patch was always at a fixed luminance of 4 cd/m^2^. In three blocks, we tested different luminances of an adapting uniform background, including 4 cd/m^2^ (“HDR gray”, no change), 40 cd/m^2^ (“HDR light gray”), and 400 cd/m^2^ (“HDR white”). We tested two additional control blocks, one consisting of a narrower SDR luminance range with a light gray adapting blank (“SDR” block), and the other approximating a classic condition with uniform background and uniform flanker orientation, with a light gray adapting blank (“Uniform” block). Blocks consisted of 400 trials each and were presented in the following order: Uniform, HDR light gray, SDR light gray, HDR gray, and HDR white.

Within each block, stimuli consisted of 45° and 135° Gabors, 4 cycles/° (i.e., the bars have a thickness of 0.125°), and 1° full width at half maximum Gaussian envelope, cropped to 1° × 1°, presented on a 5 × 5 array of 1° × 1° luminance patches. The spatial frequency of the Gabors is consistent with the stimulus preferences of single neurons in primary visual cortex with receptive fields at 3° eccentricity. The target was a contrast blend of Gabors at the two orientations, presented at the central patch, and subjects indicated via keypress the orientation of the stronger target Gabor (45° or 135°). The central patch luminance was 4 cd/m^2^, and flanker patches evenly log-linearly spanned a luminance range from 40 to 0.4 cd/m^2^ (10× to 0.1× the target patch luminance) for HDR conditions and from 12.6 to 1.26 cd/m^2^ (3.16× to 0.316× the target patch luminance) for the SDR condition. Each Gabor’s pixels spanned a range from 10× to 0.1× its flanker patch luminance, resulting in a peak contrast including Gabors of 10,000 to 1 (400 to 0.04 cd/m^2^) for the HDR array and 1000 to 1 (126 to 0.126 cd/m^2^) for the SDR array.

The target patch consisted of two Gabors (45° and 135°) at one of five possible contrast mixtures of 70%:30%, 60%:40%, 50%:50%, 40%:60%, or 30%:70%. These contrast mixtures were logarithmically applied to each full-contrast Gabor whose brightest and darkest pixels were 10 and 0.1 times its patch luminance, such that 50%:50% means that the brightest and darkest pixels of both Gabors are 3.2 and 0.32 times their patch luminance. All Gabor patterns were well above the threshold contrast visibility for normal vision. At the minimum average luminance of 0.4 cd/m^2^, in a field of view 1° × 1° and a modulation frequency of 4 cyc/°, (Barten 2003) indicates a minimum visible contrast ratio of 1.03; for a field of view 0.5° × 0.5°, the minimum visible contrast ratio of increases only to 1.06.

We tested two orthogonalized conditions to determine whether the most influential flankers were the brightest flankers, or the flankers that were most similar to the target in luminance. Each 5 × 5 grid consisted of an inner ring of 8 patches and an outer ring of 12 patches. The flankers of interest were balanced within each ring, such that there was an equal number of co-oriented and orthogonal flankers, and their locations were spatially balanced in the horizontal and vertical directions to avoid highly asymmetric patterns. In the “brightest” condition, the flanker patches of interest were the 12 brightest patches. In the “similar” luminance condition, the flanker patches of interest were the 12 patches most similar in luminance to the target patch. For both conditions, we defined “Flankers condition A” as the case in which the Gabors at the patches of interest were tilted 45° and remaining patches tilted 135°, and vice versa for “Flankers condition B”. The five target mixtures were tested in all conditions.

At the start of every trial, a black fixation cross appeared at the center of a blank screen (this adapting uniform blank screen is subsequently referred to as the “blank” screen). Participants hit the spacebar when they were ready to begin the trial. This keypress did not advance the trial unless the blank screen had been on for a minimum of 500 ms. The keypress initiated a sequence that began with the offset of the blank screen and fixation cross, replaced by a black screen (0.006055 cd/m^2^). The 5 × 5 flanker array appeared 50 ms later (three video frames; the MATLAB trial timestamp for array onset was at 39 ms, synchronized to the timestamp for blank offset), and then the target Gabors 17 ms after that (one video frame, the MATLAB trial timestamp for target onset was at 67 ms). The target Gabors remained on the screen for 250 ms and then offset, leaving the flankers on the screen until the participant hit the left of right arrow keys to report whether the stronger target Gabor was oriented to the left or the right. After keypress, the blank screen reappeared after a 500 ms wait, ending the trial.

#### 2.8.1 Experiment 2 Behavioral Analysis

Behavioral choices were fitted with a psychometric function based on a cumulative Gaussian distribution, using Psignifit software running in Python (Fründ et al. 2011). Psignifit estimates a free guessing and lapse rate parameter (options.expType = “YesNo”) to fit the behavior via maximum likelihood. Compared with classic logarithmic fitting tools, Psignifit is thought to provide better confidence intervals by avoiding two critical assumptions of stability and binomial distribution, to account for factors such as learning, fatigue, and fluctuations of attention.

To analyze the significance of the flanker-induced bias for each condition, we defined a significant behavioral effect, as cases where the 5%:95% confidence interval at the 50%-choose-“A” threshold of each curve did not cross the other curve, for the two curves from Flankers condition A and Flankers condition B. We analyzed this bias separately for the “brightest” and “similar” conditions and each of the blocks. For population analysis, we defined each subject’s bias as the difference in the two curves’ target mixtures at the 50%-choose-“A” threshold. We then applied a one-sample two-tailed t-test to examine the significance of this bias for each condition versus a null hypothesis of zero bias across the population. We also compared these biases across the “brightest” and “similar” conditions via two-tailed paired t-test. Significance tests were not corrected for multiple comparison.

#### 2.8.2 Experiment 2 EEG Data Collection and Analysis

EEG data were recorded from 64 active scalp electrodes using a Biosemi Active Two system. Electrodes were re-referenced offline prior to analysis to the average of two external electrodes placed on the left and right mastoids. Three participants had an online sampling rate of 2048 Hz while the remaining five had a sampling rate of 512 Hz. All participants’ data were down-sampled to 512 Hz offline prior to analysis. A bandpass filter (0.1 to 40 Hz) was applied offline prior to analysis.

We analyzed EEG data in Task 2 via a combination of custom MATLAB scripts and EEGLab functions (Delorme and Makeig 2004). To generate event-related potentials (ERPs), the continuous data were epoched into trials (−200 to 600 ms post blank offset or target onset depending on the analysis; see Results section). Each trial was then baseline-corrected by subtracting the average voltage in the −200- to 0-ms window from the entire waveform. We used EyeLink to track the movements of one eye (typically the right eye) at 1000 Hz. Trials in which the eye moved more than 2° during target presentation were excluded from analysis. Additionally, trials in which a given electrode’s amplitude exceeded 100 μV were also excluded. We limited the EEG analysis to trials in which the target was ambiguous (i.e., trials where the contrast mixture of the two Gabors was 40%:60%, 60%:40%, or 50%/50%). These exclusion criteria resulted in an average of 152 (*SE* = 28) usable trials per participant in the uniform block, 133 (*SE* = 26) in the HDR light gray block, 113 (*SE* = 28) in the HDR gray block, and 132 (*SE* = 21) in the HDR white block.

## 3. Results

### 3.1 Experiment 1: Dynamic Luminance Visual Acuity Task

We examined how visual acuity for letters was degraded by challenging luminance conditions. The task is a computerized variant of the standard ETDRS LogMAR test (Stewart et al. 2006), in which subjects verbally report the identities of five letters appearing on each trial, with decreasing letter sizes across trials. The difference in our task was that we added a preceding background flash and modified the luminance and stimulus duration parameters to approximate the conditions of the subsequent target discrimination task (Task 2) to provide a comparison of the HDR target discrimination task and standard visual acuity tasks. Results of Task 1’s acuity measurements are shown in Fig. 5. The vertical axis shows acuity expressed as the LogMAR, and as the equivalent Snellen ratio, which shows the resolution of the test participant’s vision at 20 ft (numerator value) compared with the distance at which a person with normal vision would have the same line resolution ability (denominator). With normal vision, a Snellen score of 20/20, the minimum angle of resolution is 1 arcmin, corresponding to a LogMAR score of zero.

**Fig. 5.**
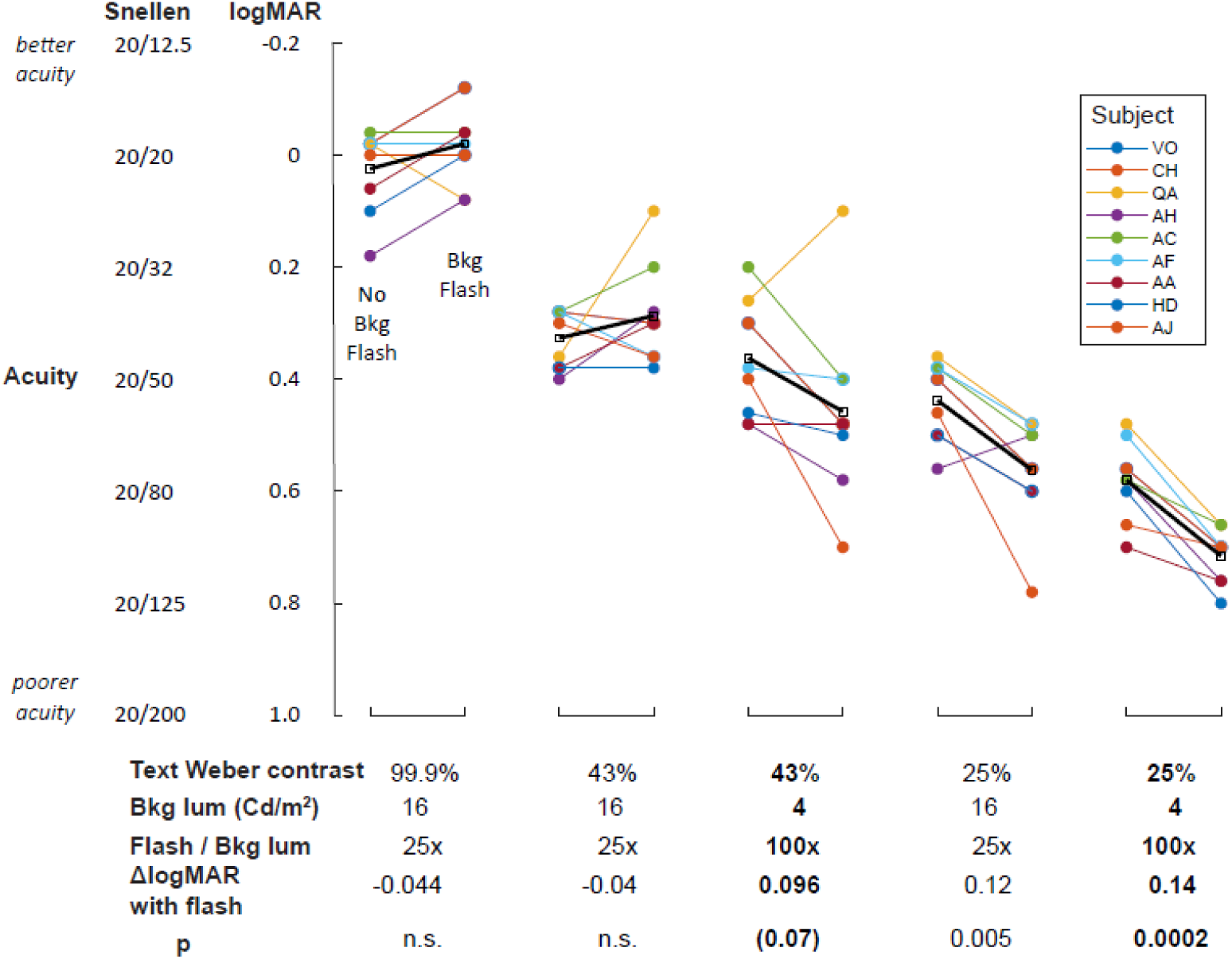
Dependence of visual acuity on flash, background luminance, and text contrast (Task 1); N = 9 subjects

As expected, acuity generally decreased as text contrast and background luminance decreased, but the details of the decrease depended on the parameters. When a background flash of 400 cd/m^2^ (25× and 100× the letter background for 16 and 4 cd/m^2^ backgrounds, respectively) preceded each letter onset, some test subjects gained acuity and some test subjects lost acuity, but the average changed little.

At a text Weber contrast of 99.9% and a steady background luminance of 16 cd/m^2^, the average acuity of test subjects was 0.02 LogMAR. When preceded by a background flash, average acuity improved nonsignificantly to 0 LogMAR. Decreasing the text contrast from 99.9% to 43%, while holding the background luminance at 16 cd/m^2^, acuity decreased to 0.35 LogMAR without flash and to 0.31 LogMAR with flash. At a constant text contrast of 43% and further decrease in background luminance to 4 cd/m^2^, average acuity was 0.38 LogMAR without flash and 0.48 LogMAR with flash—a nearly significant (p = 0.07) degradation of 0.1 LogMAR. At further reduced text contrast of 25%, with background luminances of 16 and 4 cd/m^2^, the addition of a flash significantly decreased acuity by 0.12 and 0.14 LogMAR (p = 0.005 and 0.0002), respectively, to 0.58 and 0.70 LogMAR, corresponding to Snellen ratios of approximately 20/80 and 20/125.

Overall, these results show that the visual system is able to rapidly normalize for changes in luminance, such that flashes have negligible impact on letter acuity within 200 ms immediately following the flash, for flashes up to 25× the background luminance and text Weber contrast above 43%. At more challenging conditions of flashes 100× the background luminance and/or weaker text contrast of 25%, flashes induced a mild acuity loss of up to 0.14 LogMAR. However, even at the most challenging condition, the resulting acuity of 0.70 LogMAR indicates that the minimum angle of resolution is 5 arcmin (0.083°), well within the 0.125° receptive field size of simple cells in V1 and below the width of Gabor bands in our Task 2 stimuli. Thus, although flashes at 100× the background luminance can reduce visual acuity, sufficient acuity should remain to perform the Task 2 target discrimination task and to observe any luminance-related contextual effects.

### 3.2 Experiment 2: HDR Luminance Target Discrimination Task

Previous reports of contextual effects on target orientation discrimination were based on uniform flanker lines against a uniform, static background luminance. In the simplified classic example in Fig. 6A, a low-contrast central Gabor target is surrounded by two flankers at higher contrast, and the flankers are either co-aligned (left), or rotated orthogonally to the target (right). In this example, the right target is more visible than the left target because the left target’s visibility is suppressed by the surrounding co-aligned flankers (assimilation). Conversely, at even lower target contrast (not shown), the left target becomes more visible than the right target because of facilitation by the co-aligned flankers. In EEG recordings, flanker co-linearity also produced an increased midline occipital positive polarity between 80 to 140 ms after stimulus onset, consistent with a mechanism in Area V1 (Polat and Norcia 1996; Khoe et al. 2004). This dichotomy of contextual facilitation versus suppression to static SDR stimuli has formed the basis of computational models of V1, based on a balance of recurrent excitation and inhibition (Chen and Tyler 2002; Li 2011).

**Fig. 6.**
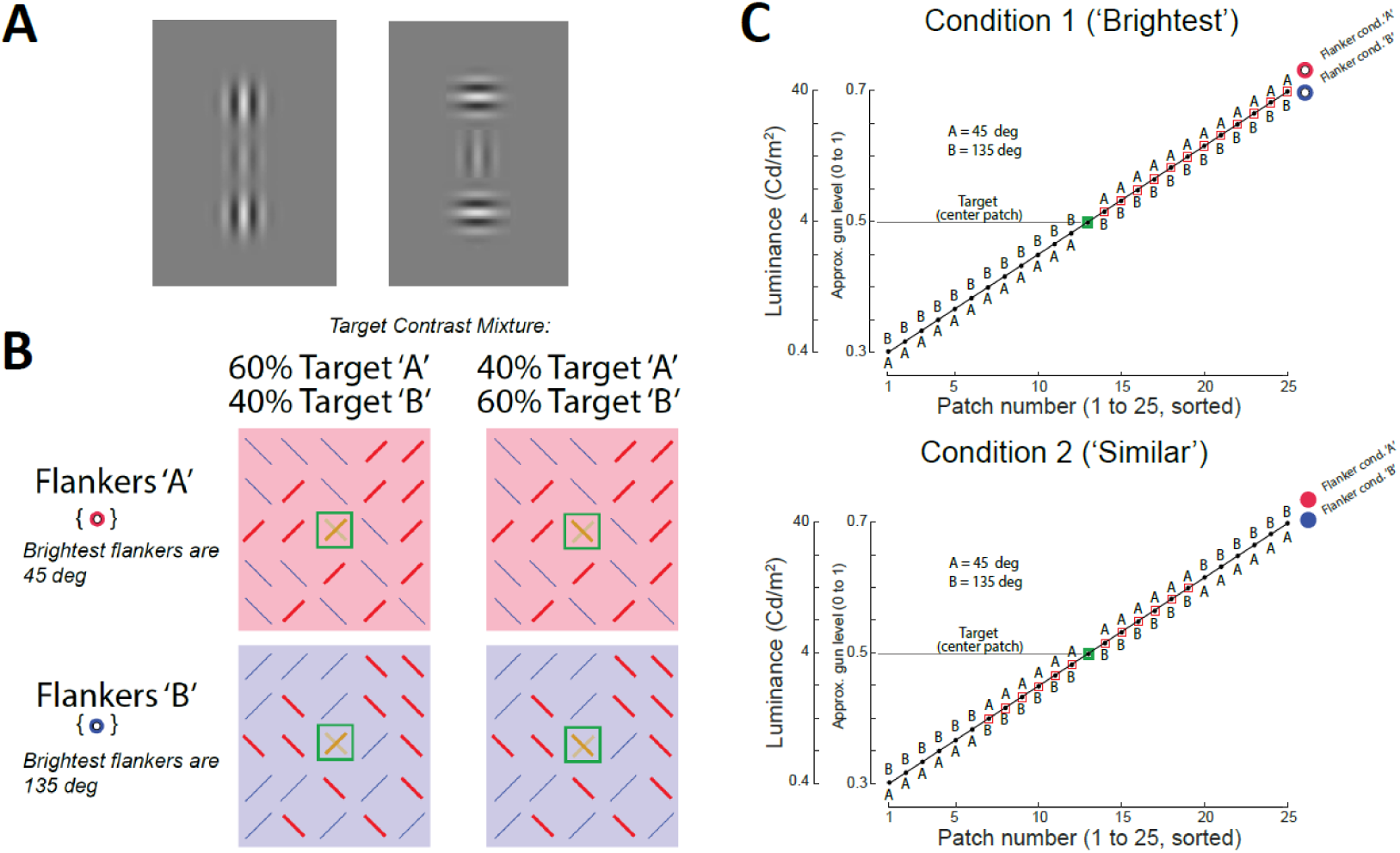
Stimulus design for HDR luminance target orientation discrimination task (Task 2). (A) Example of classic suppression/assimilation effect when the central target is high contrast. The target in the left image is less visible because it is co-oriented with its flankers, whereas the target in the right image is more salient because it is orthogonal to its flankers. (B) Schematic of example trial types for “Brightest condition” for Example 1. The brightest flankers (red lines) are oriented either 45° (Flankers “A”) or 135° (Flankers “B”). Only two of five possible target contrast mixtures are shown. For the “Similar” condition (not shown), the flankers that are most similar in luminance to the target would be highlighted in red and oriented either 45° (Flankers “A”) or 135° (Flankers “B”). C) Luminance distributions of target and 24 flankers. Flanker patches span 0.4 to 40 cd/m^2^ (0.04 to 400 cd/m^2^ full range including Gabors).

To understand and model contextual mechanisms of luminance normalization under real world luminance dynamics, we introduced two variations to the classic flanker task: 1) a preceding adapting background to mimic the luminance change across gaze shifts, and 2) a 5 × 5 array of luminance patches, spanning a 10- or 100-fold difference in luminance, to mimic the conjunction of form and luminance in naturalistic scenes. This combination of adapting background, patches, and Gabors resulted in a total luminance range of up to 10,000 to 1.

We tested this combination via a two-alternative forced choice task in which subjects report the orientation of the stronger of two Gabor targets shown at the center of the 5 × 5 array (Fig. 2B). By fitting the behavioral responses across target contrast mixtures with a psychometric function, we were able to determine whether the flankers induced a facilitatory or suppressive/assimilation effect under real-world luminance dynamics. Additionally, by manipulating the conjunction of patch luminance and patch orientation via two orthogonalized conditions, we were able to test alternative hypotheses about normalization mechanisms that predict whether the brightest flankers, or the flankers that have most similar luminance to the target, would have stronger effect.

To examine how contextual luminance and orientation combine to affect target discrimination, we manipulated the conjunction of luminance and orientation across the 5 × 5 array of patches. Fig. 6B illustrates schematic examples of such stimuli for the “brightest” condition in which the flankers of interest (indicated by red lines) are at the brightest patches. In the upper left example of Fig. 6B, corresponding to the stimulus example in Fig. 2B, the target mixture is 60%:40% (one of five possible target mixtures, 30%:70%, 40%:60%, 50%:50%, 60%:40%, 70%:30% A:B) and the Flanker condition is “A”, so that both the target and the flankers of interest are tilted to the right (45°). The comparison condition with the identical target mixture is Flanker condition “B” (lower left), in which the flankers of interest are tilted to the left (135°). Fig. 6C (top) illustrates this comparison of Flankers A (red open circle) versus Flankers B (blue open circle) orientations for the “brightest” condition, sorted by patch luminance. The “A”s indicate 45° flankers, the “B”s indicate 135° flankers, and the red squares indicate the flankers of interest (the brightest patches, in this condition). If the flanker effect is driven by the brightest flankers, we would expect to see a difference in the target report between these two conditions Flankers A versus Flankers B in the “brightest” condition. The same direction of bias should be present across a range of target mixtures, including when the target mixture is 40%:60% (upper and lower right examples in Fig. 6B).

Conversely, if the flanker effect is driven by the flankers that are most similar in luminance to the target, the effect would cancel out in the “brightest” condition (flankers near the target luminance are both co-oriented and orthogonal). The flanker effect would only be observed in the “similar” condition, in which the flankers of interest are at the 12 patches most similar to the target in luminance (Fig. 6C bottom).

An example of the behavioral results is shown for one subject “VO” in Fig. 7. As expected, the subject tended to choose “A” (45° target) when the target contrast mixture was 60%:40% or 70%:30%, and choose “B” (135° target) when the target contrast mixture was 30%:70% or 40%:60%, indicating that the subject is generally able to correctly perceive the target’s dominant orientation. The psychometric function did not significantly differ for the Flankers A (red circles) and Flankers B conditions (blue circles) for the “brightest” condition (Fig. 7A, B, and C), across all three adapting blank luminance levels 400, 40, and 4 cd/m^2^, indicating that the brightest flankers failed to bias behavioral choice.

**Fig. 7.**
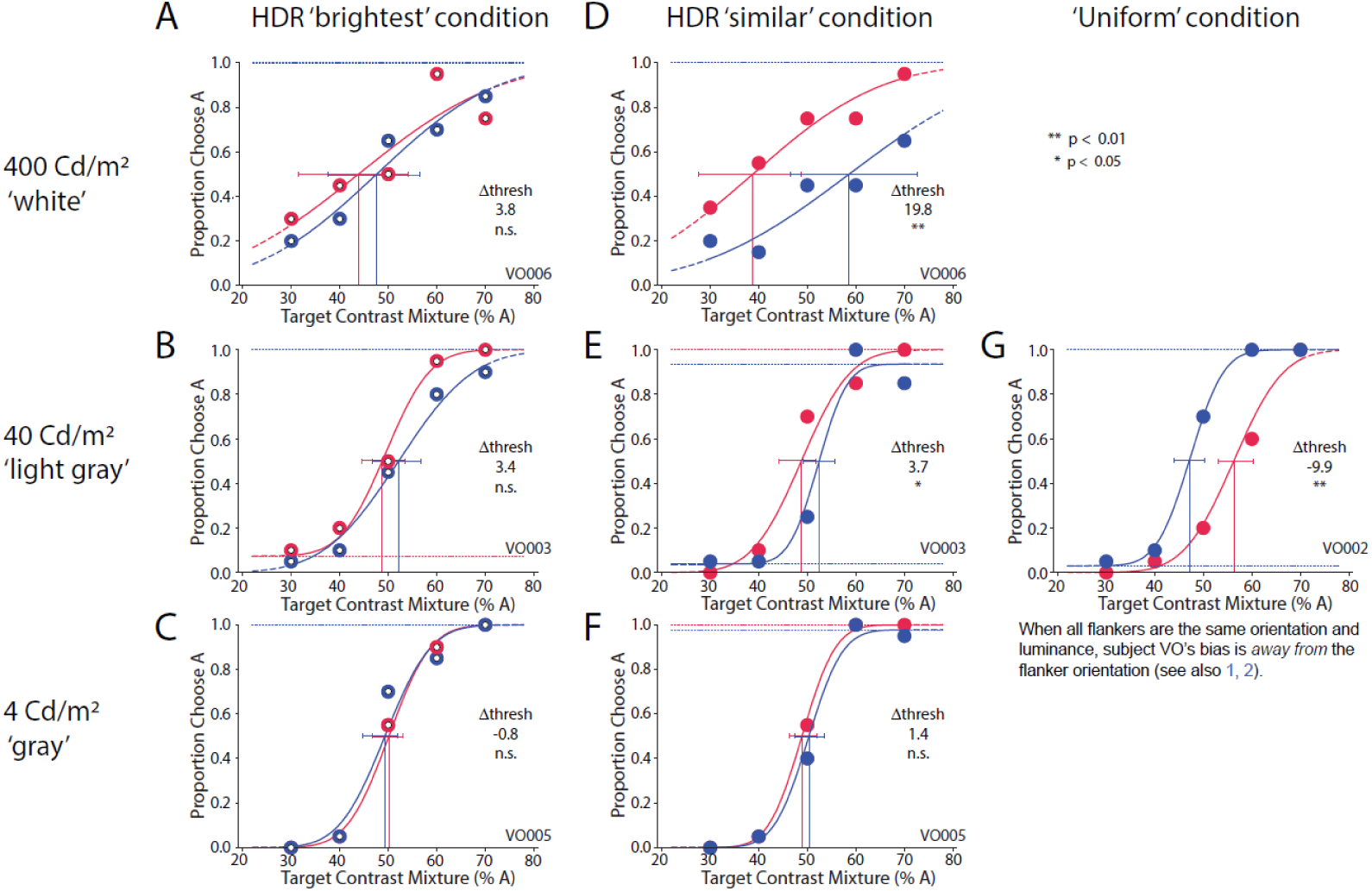
Subject VO’s behavioral results on reporting the orientation of the target Gabor broken out by block and condition. Red lines and data refer to thresholds when the flanker Gabors of interest (the brightest flankers for panels A–C, or the flankers most similar to the target in luminance for panels D-F) are tilted to the right (45°, Flankers “A”); blue lines and data refer to thresholds when the flanker Gabors of interest are tilted to the left (135°, Flankers “B”). Error bars show 5%–95% confidence intervals based on Psignifit.

Conversely, the flankers induced a significant bias on target choice behavior in the “similar” condition, when the flankers of interest were at the 12 patches most similar in luminance to the target. The effect was facilitatory; there was a bias in the subject’s response toward the orientation of the flankers of interest, as indicated by the leftward shift of the red curve (increased likelihood to choose “A” when the flankers of interest are 45°) and the rightward shift of the blue curve (increased likelihood to choose “B” when the flankers of interest are 135°) in Fig. 7D and 7E. This flanker-induced bias was significant for the two brightest adapting screen luminances, 400 and 40 cd/m^2^ (p < 0.01 and p < 0.05, respectively), but it was not significant for the lowest adapting screen luminance, 4 cd/m^2^ (i.e., where the luminance at the target patch was unchanged [Fig. 7F]). In the “Uniform” condition (Fig. 7G), when all flankers had the same orientation and the same 4-cd/m^2^ patch luminance with a 40-cd/m^2^ adapting blank, the subject’s report was significantly biased away from the flanker orientation, consistent with suppression or assimilation (p < 0.01).

Another subject, AH, also showed a flanker-induced bias toward facilitation that was significant in both the “brightest” and “similar” conditions for the 400-cd/m^2^ “HDR white” adapting blank (Fig. 8A and 8D; p < 0.01 in both cases). The strength of this threshold bias, a difference of 35.5% target contrast mixture at the 50%-choose-“A” threshold in the “brightest” condition, is illustrated by the fact that even when the target mixture is 70% “B” (30% “A”), the subject chose “A” 60% of the time under Flanker condition “A”, and the subject chose “B” 97% of the time under Flanker condition “B”. Conversely, when the target was 70% “A”, the subject chose “B” 45% of the time (chose “A” 55% of the time) under Flanker condition “B”, and chose “A” 90% of the time under Flanker condition “A”. As with subject VO, subject AH also showed a facilitatory bias for the “similar” condition with the 40 cd/m^2^ “HDR light gray” adapting blank (Fig. 8E) and no significant bias in the remaining HDR conditions (Fig. 8B, 8C, and 8F). However, unlike subject VO, subject AH showed no bias in the “Uniform” condition.

**Fig. 8.**
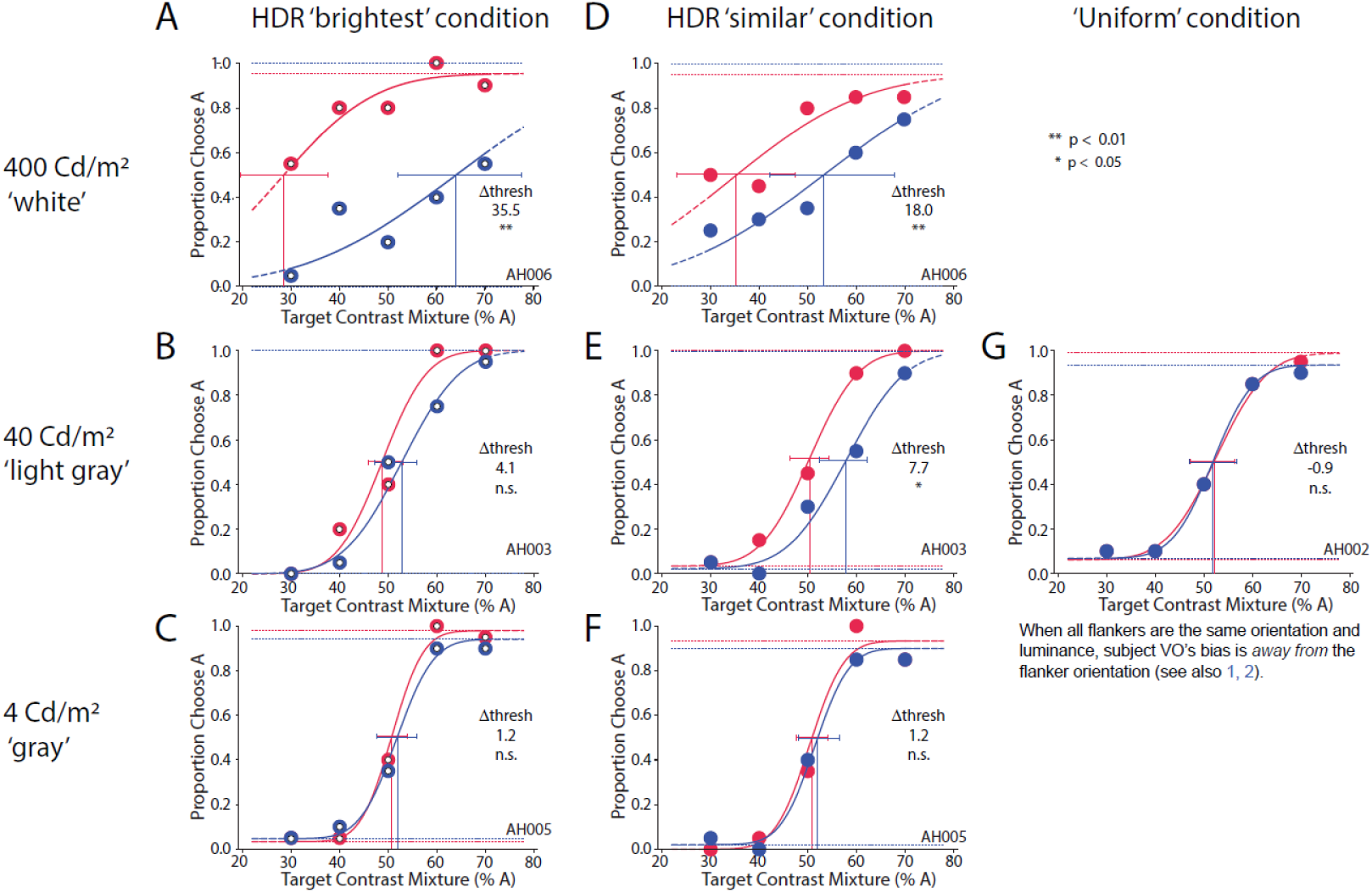
Subject AH’s behavioral results on reporting the orientation of the target Gabor broken out by block and condition. Red lines and data refer to conditions when the flanker Gabors of interest are tilted to the right (45°, Flankers “B”); blue lines and data refer to conditions when the flanker Gabors of interest are tilted to the left (135°, Flankers “B”).

Across subjects, there was a strong flanker-induced bias toward facilitation when the adapting screen was much brighter than the Gabor display (i.e., there was a strong bias to report targets as co-oriented with the flankers of interest) (Fig. 9). At the brightest adapting blank of 400 cd/m^2^ (100× brighter than the target patch, HDR white”, this flanker-induced facilitation was strong and significant for both the “brightest” and “similar” conditions (threshold bias = 12.0 ± 12.1 and 14.8 ± 7.2, p = 0.018 and p = 0.0003, respectively), and there was no significant difference between these two conditions, indicating that both the brightest flankers and the flankers similar to the target in luminance contributed to the facilitation.

**Fig. 9.**
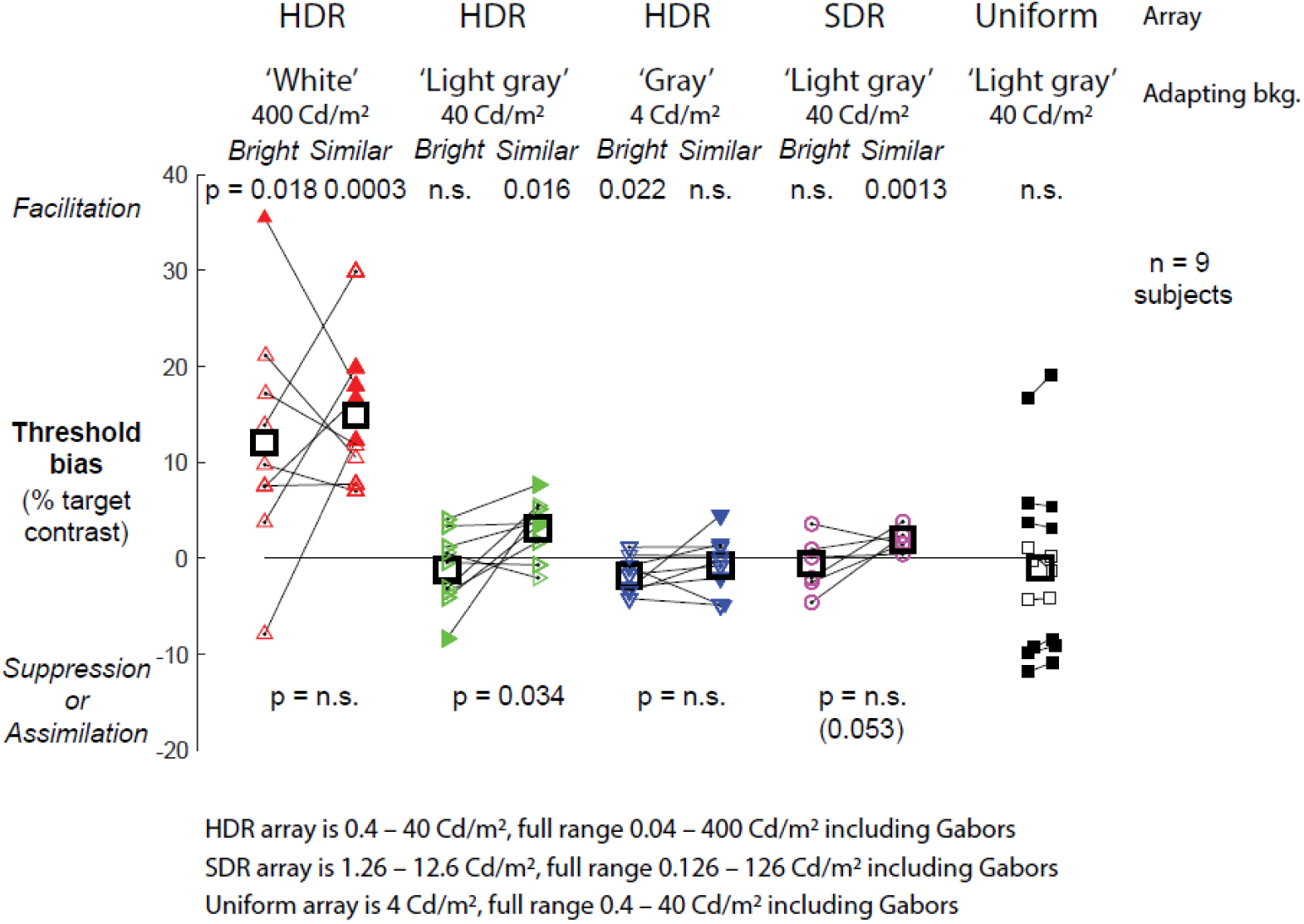
Population results for Gabor orientation discrimination. For the Uniform condition, connected squares in left and right columns show results of split trials, indicating the precision of the measurement for each subject.

The effect was titrated by the magnitude of the luminance change. When the adapting screen was only 10× brighter than the target patch (“HDR light gray”), the flanker-induced facilitation was significant for the “similar” condition (threshold bias = 3.1 ± 3.1, p = 0.016), but this bias was abolished in the “brightest” condition, indicating that it was not driven by the brightest flankers (−1.2 ± 4.0, p = not significant [n.s.]). This difference between “similar” and “brightest” conditions was significant (p = 0.034, two-tailed paired t-test).

When the adapting screen was the same luminance as the target patch (“HDR gray”), there was a weak but significant bias toward suppression in the “brightest” condition (−1.7 ± 1.8, p = 0.022), consistent with previous reports of suppression to high-contrast targets at static luminance. This bias was abolished in the “similar” condition (−0.06 ± 3.0, p = n.s.), but the difference between conditions was not significant.

Could these effects have been observed with an SDR display? We tested an “SDR light gray” condition in which the array patch luminances spanned only 12.6 to 1.26 cd/m^2^ (3.16× to 0.316× the target patch luminance, a 10× range), versus 40 to 0.4 cd/m^2^ (10× to 0.1× the target patch luminance, a 100× range) in the HDR array. As with the “HDR light gray” condition, there was a significant flanker-induced facilitation in the “similar” condition (1.9 ± 1.0, p = 0.0013), but this effect was abolished in the “brightest” condition (−0.5 ± 2.5, p = n.s.). The difference between these two conditions approached significance (p = 0.053). Thus, the effect was much weaker under SDR but consistent with the results under HDR.

Across subjects, the “uniform” condition resulted in a wide variation of individual biases (−1.0 ± 9.4), ranging from significant facilitation (18.0, p < 0.01) to significant suppression (−11.8, p < 0.01). The bias was not due to noise in the measurement, as randomly splitting the trials resulted in almost no change to individual biases (Fig. 9, pairs of connected squares). The direction of this individual bias did not appear to be related to the magnitude of facilitation in the HDR and SDR conditions. The wide range of individual biases under the “uniform” condition contrasts with the narrower range of biases in the HDR conditions, and especially the SDR light gray “similar” condition (1.9 ± 1.0). It also contrasts with previous reports of target suppression to static uniform flankers, indicating that our addition of the adapting light gray blank has substantially altered the conditions to produce unexpected behavior. We speculate that the blank, although only 10× brighter than the target patch and the same amplitude as the flanker Gabors, may increase ambiguity and enable the emergence of strong priors, based on the subjects’ false expectation of a relationship between the target and flankers.

Only data from the HDR white, HDR light gray, HDR gray, and uniform conditions were used in the EEG analysis. The grand average ERP waveforms, time-locked to the offset of the blank screen, are shown in Fig. 10. Upon visual inspection the initial positive deflection post blank offset showed graded amplitude and latency depending on the magnitude of luminance decrement after the blank, with largest amplitude and shortest latency for the largest luminance change. To assess the significance of this difference, we used a leave-one-out jackknife analysis method (Ulrich and Miller 2001). We measured peak amplitude as the maximum amplitude of the waveform within a 0- to 150-ms time window post blank offset, and the latency was the time at which the component reached 50% of that peak amplitude. A repeated measures analysis of variance (ANOVA), with corrected F-value to account for the decreased variability in the jackknife method, found no significant difference between the peak amplitudes across conditions, *F*(3,7) = 0.34, *p* = 0.79. However there was a significant difference in peak latency across conditions, *F*(3,7) = 16.56, *p* < 0.001.

**Fig. 10.**
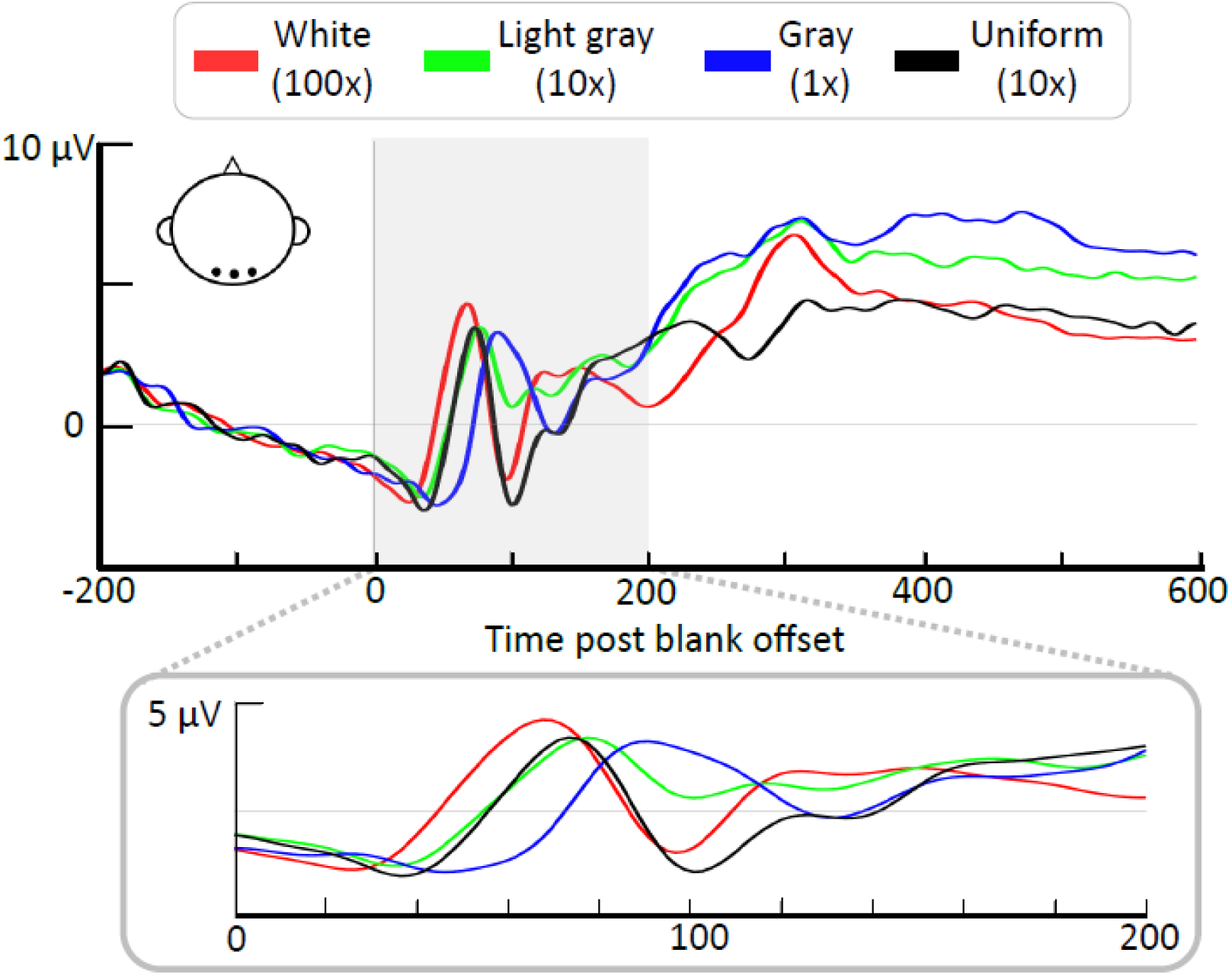
Top: Grand average waveforms for the uniform and HDR blocks (luminance difference included in parentheses) calculated from early visual area electrodes (O1, O2, and Oz). Note that waveforms here are time locked to the blank screen offset. Bottom: Same graph as top but with the 0- to 200-ms post blank screen offset time window expanded.

The critical comparison, however, was between trials where participants reported the target as having an orientation that was congruent or incongruent with the background flankers (Fig. 11). To analyze this difference, epochs time-locked to the onset of the target were used, and a difference waveform was calculated by subtracting the incongruent waveform from the congruent waveform. Previous work using a similar subtraction method for static uniform flankers had found a significant difference in the 80- to 150-ms window post target onset (Khoe et al. 2004). Based on those results, we analyzed the same time window to test for a difference between conditions. We used the same jackknife method with a corrected F-value, repeated measures ANOVA as described previously. We observed no significant flanker effect on ERP amplitude in any condition, whether pooled across “brightest” and “similar” conditions or analyzed separately, and no significant difference between conditions, *F*(3,7) = 0.93, *p* = 0.44.

**Fig. 11.**
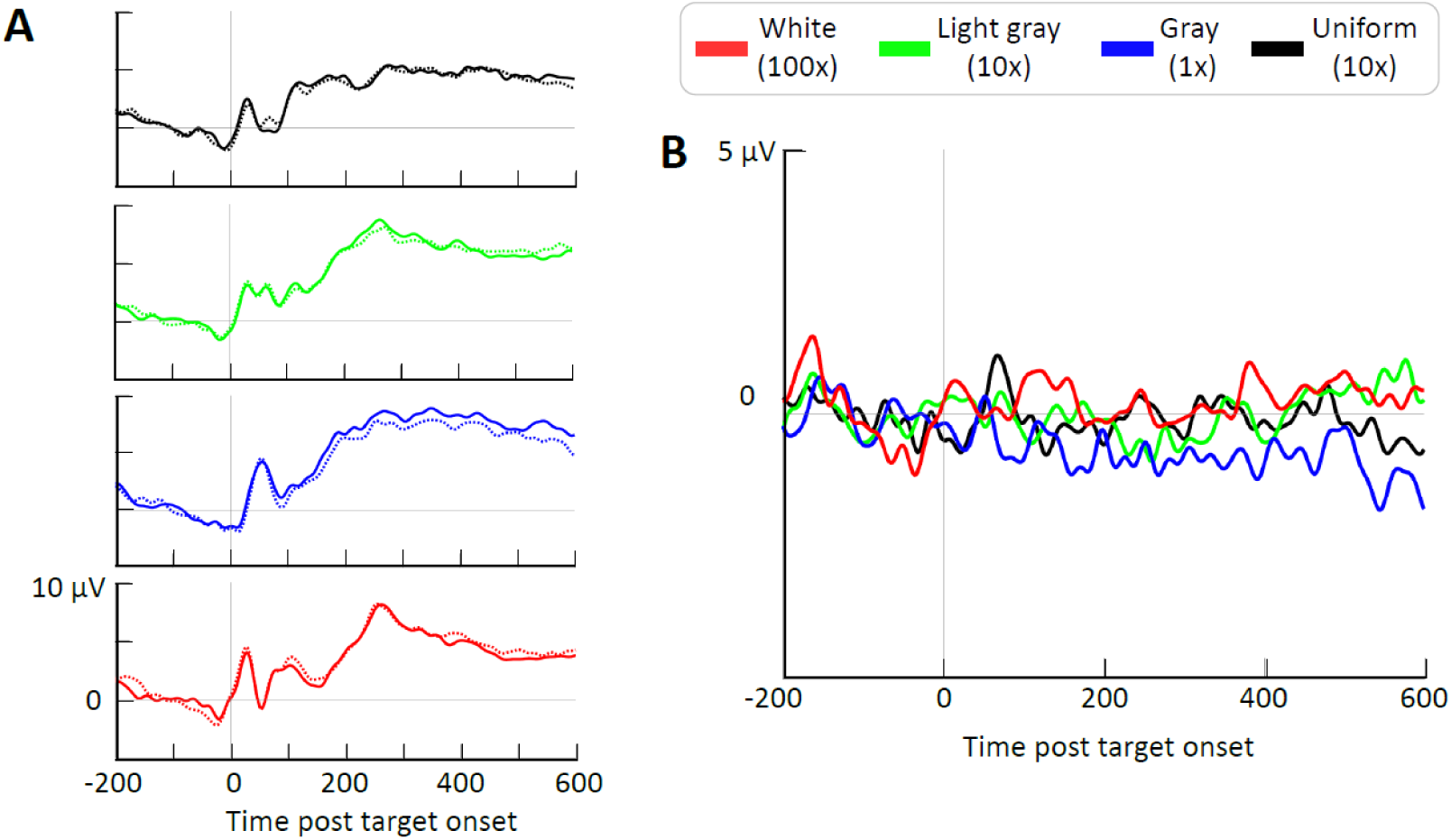
A) Congruent (dotted lines) and incongruent (solid lines) plotted for the uniform (black), HDR light gray (green), HDR gray (blue), and HDR white (red) blocks. B) Difference waves (congruent minus incongruent) for each block.

## 4. Conclusion

These results advance our capability to develop resilient and intuitive real-world machine vision, by discovering HDR luminance normalization mechanisms associated with primary visual cortex. We developed an HDR display research platform with improved characteristics of more than 100,000-to-1 contrast ratio and pseudo 11 bits, versus standard SDR displays that are typically limited to 1000-to-1 contrast ratio and 8–10 bits. This advance allowed us to discover new phenomena linking contextual mechanisms to luminance normalization and target discrimination. We showed for the first time that abrupt darkening (as would occur during gaze shifts) induces facilitation, even for high-contrast targets. The effect was titrated by the magnitude of the luminance change. Surprisingly, the effect was driven by flankers with similar luminance to the target, whereas classic mechanisms such as divisive or subtractive normalization predict that the effect should be driven by the brightest flankers. We also showed that the effect required an HDR stimulus to be observed, and was much weaker (but significant) under SDR. We showed that the classic case of suppression by uniform flankers becomes surprisingly ambiguous, with large individual variations toward both facilitation and suppression, with the simple addition of a 10× darkening. We showed that such large luminance changes manifest as shorter latency ERP deflections, although we were unable to observe differences in ERP amplitude related to flanker co-orientation. In addition we showed that letter acuity is not substantially altered by a preceding flash, except under the most challenging conditions, and we tied this back to the standard LogMAR test of visual acuity. Together, these results provide the framework necessary to construct computational vision models that leverage biology’s substantial advantages in handling high-dynamic range images.

## 5. Discussion

An ongoing challenge to Army modernization is how to develop autonomous teammates that can function in the real world for effective teamwork. Recent advances in machine vision, based on deep neural networks (DNNs) trained on large SDR photographic and synthetic databases, have resulted in substantial improvements in automatic target recognition capability, but a substantial problem remains of unexpected misclassifications, which limit machine vision credibility and require too many user interventions to be practicable. For example, a recent report showed that DNNs trained on ImageNet photographs are over-reliant on texture for object classification, and are easily fooled by synthetic images in which object surfaces are replaced by other textures (Baker et al. 2018; Geirhos et al. 2018). Whereas biological vision has many mid-level processes to support resilient generalization, such processes are absent in DNNs. This is evidenced by the limited ability of DNNs to explain brain activity in many visual areas and by the ongoing challenge to resolve the disjunction between machine versus human patterns of classification errors. A strategy to resolve this capability gap is to incorporate biological resilience into machine vision.

Our work on HDR luminance normalization aligns with this strategy by building on biology’s advantage in normalizing HDR images, with the aim of building tone-mapping and saliency models that can feed into DNNs to improve overall performance. We took the approach of focusing on contextual processing in V1, which has been well explored for the case of static SDR images and is supported by a large body of work in human and animal behavior, anatomy, and physiology, and we advanced it toward real-world application by adding HDR luminance dynamics consistent with gaze shifts in naturalistic scenes. We report unexpectedly strong behavioral results that show contextual facilitation following abrupt darkening, even for high-contrast targets, and an unexpected phenomenon of contextual luminance-similarity-dependent facilitation that is consistent with traditional Gestalt theories of grouping. Both of these results challenge models based on recurrent excitation and inhibition that would predict stronger effects from the brightest flankers. Our future work will be to incorporate these results into computational modeling, based on both recurrent excitation/inhibition modeling and on saliency modeling.

The behavioral results in the target discrimination task highlight limitations of classic laboratory-based approaches to studying biological vision, namely the limited generalizability to real-world dynamics. By simply widening the luminance range and adding an abrupt luminance change, we observed novel phenomena (facilitation to high-contrast targets, and flanker effects driven by luminance similarity to the target) that are not predicted by previous models of contextual effects. Under the HDR gray condition (no luminance change), we reproduced the classic suppression effect. However, the flanker effect became highly variable across subjects in the uniform condition in which we added a novel 10× darkening, leading to the unexpected observation of both facilitation and suppression across different subjects to identical stimuli. This supports the inherent ambiguity of vision and the role of top-down prior expectations or beliefs on individual variability in behavioral performance.

EEG data were analyzed to pinpoint when in the visual processing pipeline the effect of the luminance change caused the behavioral effect seen here. The EEG components time-locked to the offset of the blank screen showed the expected trend of earlier onset latency when the luminance change between the blank and array screen was larger. This result was expected, as the greater the luminance change, the more salient the transition. Increasing saliency of visual input has previously been shown to elicit earlier onset in components linked to attentional processes (Töllner et al. 2011). Our data did not, however, replicate previous findings that target-flanker orientation congruency modulates the ERP amplitude (Khoe et al. 2004). We found no such congruency effect on ERP amplitude in our task. This difference in results is perhaps unsurprising given the differences in the task used in previous work and that used here. One such difference is that in the previous study, Khoe and colleagues only presented participants with two flanker Gabors of the same orientation. By contrast, our stimuli had a dense array of Gabors in both congruent and orthogonally incongruent orientations, to optimize for behavioral analysis. This increase in complexity of the visual input may have introduced more noise into the EEG data, data known to have a low signal-to-noise ratio (Luck 2014), and therefore made the signal difficult to detect through traditional analyses. Moreover, the task was very different in this experiment compared with the previous study. Previously a detection task was used, where participants had to report whether a Gabor was presented. In the current work, participants are tasked with determining which of two Gabors has the higher contrast. This may bring in an additional decision-making component to the task that is not there when participants are only required to report the presence or absence of stimulus.

The generalizability of these results is limited by several factors. First, our display range of 100,000 to 1 is still well below the 10^9^-to-1 range of some natural scenes. However, scenes that exceed 100,000-to-1 luminance range are likely to span very wide fields-of-view and may involve staring at a light source such as the sun, and a display with such capability could be damaging to the eye. By capturing a substantial portion of both mesopic and photopic ranges, our display captures the key transition between indoor and outdoor illumination that poses some of the most common HDR luminance challenges. Another limitation is that in our target discrimination task, the subject maintained fixation while the luminance changed, whereas such luminance changes typically occur across gaze shifts in a static scene. An important difference is that in gaze shifts the visual system has presaccadic information about the luminances at the target postsaccade. It is unknown whether this would alter our behavioral results, so a future plan is to repeat this experiment under controlled free-viewing, where the subject shifts his/her gaze from bright to dark regions in a static image. Finally, another factor limiting the generalizability of these results is that these experiments were conducted with individual test subjects, whereas optimal teaming behavior may require that the machine vision encompass both human capabilities and super-human capabilities. Depending on constraints such as size, weight, and processing power, it may not be optimal from a teaming framework for the machine vision to exactly match biological vision. However, we would argue that in most expected use cases of autonomy, where the drone or vehicle is mostly autonomous at the front line, reducing the frequency of required user interventions means that the largest sources of misclassification failure should be rooted out. A common denominator of many failures is improper normalization of the visual input, so tackling this issue, while taking advantage of DNN advances for later processing, efficiently leverages ongoing advances in biological and machine vision.

## Acknowledgments

We acknowledge the generous support of Dr Nigel Davies and Mario Kleiner, who provided advice and code for the LogMAR and high dynamic range display, respectively. Research was sponsored by the CCDC Army Research Laboratory (ARL) and was accomplished under CAST 076910227001, ARL-74A-HR53, and Oak Ridge Associated Universities cooperative agreement W911NF-16-2-0008. The views and conclusions contained in this document are those of the authors and should not be interpreted as representing the official policies, either expressed or implied, of the ARL or US government. The US government is authorized to reproduce and distribute reprints for government purposes notwithstanding any copyright notation herein.

## List of Symbols, Abbreviations, and Acronyms

2-D: two-dimensional
3-D: three-dimensional
ANOVA: analysis of variance
arcmin: minutes of arc
ATR: automatic target recognition
bpc: bits per channel
DNN: deep neural network
EEG: electroencephalography
ERP: event-related potential
ETDRS: Early Treatment Diabetic Retinopathy Study
HDR: high dynamic range
IR: infrared
LCD: liquid crystal display
LogMAR: Logarithm of the minimum-angle-of-resolution
n.s.: not significant
RGB: red-green-blue
SDR: standard dynamic range
V1: primary visual cortex

## Distribution List

1 (PDF) DEFENSE TECHNICAL INFORMATION CTR DTIC OCA

1 (PDF) GOVT PRINTG OFC A MALHOTRA

1 (PDF) CCDC ARL FCDD RLH FC C HUNG

## References

Adelson EH. Lightness perception and lightness illusions. In: Gazzaniga M, editor. The new cognitive neurosciences. 2nd ed. Cambridge (MA): MIT Press; 2000. p. 339–351.

Allred SR, Radonjic A, Gilchrist AL, Brainard DH. Lightness perception in high dynamic range images: local and remote luminance effects. J Vis. 2012;12(2). doi:10.1167/12.2.7

Anderson BL. A theory of illusory lightness and transparency in monocular and binocular images: the role of contour junctions. Perception. 1997;26(4):419–453. doi:10.1068/p260419.

Bach M. 135 Visual phenomena and optical illusions [accessed 2019 July 12]. https://michaelbach.de/ot/index.html.

Baker N, Lu H, Erlikhman G, and Kellman PJ. Deep convolutional networks do not classify based on global object shape. PLoS Computational Biology. 2018;14(12):e1006613.

Barten PG. Formula for the contrast sensitivity of the human eye. Proceedings of Image Quality and System Performance; 2003. doi.org/10.1117/12.5574676.

Blakeslee B, McCourt ME. A unified theory of brightness contrast and assimilation incorporating oriented multiscale spatial filtering and contrast normalization. Vision Research. 2004;44(21):2483–2503.

Borji A. Saliency prediction in the deep learning era: an empirical investigation. 2018. arXiv preprint arXiv:1810.03716.

Brémond R, Petit J, and Tarel JP. Saliency maps of high dynamic range images. In: European Conference on Computer Vision Berlin, Heidelberg (Germany): Springer; 2010. p. 118–130.

Chen C-C, Tyler CW. Lateral sensitivity modulation explains the flanker effect in contrast discrimination. Proceedings of the Royal Society of London. Series B: Biological Sciences. 2001;268(1466):509–516.

Chen C-C, Tyler CW. Lateral modulation of contrast discrimination: flanker orientation effects. Journal of Vision. 2002;2(6):8.

Chen C-C, Kasamatsu T, Polat U, Norcia AM. Contrast response characteristics of long-range lateral interactions in cat striate cortex. Neuroreport. 2001;12(4):655–661.

Clavagnier S, Falchier A, Kennedy H. Long-distance feedback projections to area V1: implications for multisensory integration, spatial awareness, and visual consciousness. Cognitive, Affective, and Behavioral Neuroscience. 2004;4(2):117–126.

Conway BR, Moeller S, Tsao DY. Specialized color modules in macaque extrastriate cortex. Neuron. 2007;6(3):560–573. doi:10.1016/j.neuron.2007.10.008.

Delorme A, Makeig S. EEGLAB: an open source toolbox for analysis of single-trial EEG dynamics including independent component analysis. Journal of Neuroscience Methods. 2004;134(1):9–21.

Foster DH, Amano K. Hyperspectral imaging in color vision research: tutorial. Journal of the Optical Society of America a-Optics Image Science and Vision. 2019;36(4):606–627. doi:10.1364/Josaa.36.000606.

Fründ I, Haenel NV, Wichmann FA. Inference for psychometric functions in the presence of nonstationary behavior. Journal of Vision. 2011;11(6):16.

Geirhos R, Rubisch P, Michaelis C, Bethge M, Wichmann FA, Brendel W. ImageNet-trained CNNs are biased towards texture; increasing shape bias improves accuracy and robustness; 2018. arXiv preprint arXiv:1811.12231.

Gilchrist A, Kossyfidis C, Bonato F, Agostini T, Cataliotti J, Li X, Economou E. An anchoring theory of lightness perception. Psychol Rev. 1999:106(4):795–834.

Hung CP, Ramsden BM, Roe AW. A functional circuitry for edge-induced brightness perception. Nat Neurosci. 2007;10(9):1185–1190.

Hung CP, Ramsden BM, Roe AW. Inherent biases in spontaneous cortical dynamics. In: Ding M, Glanzman HD, editors. Neuronal variability and its functional significance. London (UK): Oxford University Press; 2010.

Janssen P, Vogels R, Liu Y, Orban GA. Macaque inferior temporal neurons are selective for three-dimensional boundaries and surfaces. The Journal of Neuroscience: the Official Journal of the Society for Neuroscience. 2001;21(23):9419–9429.

Jiang X, Bollich A, Cox P, Hyder E, James J, Gowani SA, Riesenhuber M. A quantitative link between face discrimination deficits and neuronal selectivity for faces in autism. NeuroImage: Clinical. 2013;2:320–331. http://dx.doi.org/10.1016/j.nicl.2013.02.002.

Keezer MR, Pelletier A, Stechysin B, Veilleux M, Jetté N, Wolfson C. The diagnostic test accuracy of a screening questionnaire and algorithm in the identification of adults with epilepsy. Epilepsia. 2014;55(11):1763–1771.

Khoe W, Freeman E, Woldorff M, Mangun GR. Electrophysiological correlates of lateral interactions in human visual cortex. Vision Research. 2004;44(14):1659–1673.

Kleiner M. Psychtoolbox software. Personal communication; 2017. Newer generation cards (e.g., the AMD Volcanic Islands series) are currently software limited to at most 11 bpc due to the current graphics driver design. A special xorg.conf setup is needed to enable this experimental mode, as described in the “help PsychImaging” section for ‘EnableNative16BitFramebuffer’, and it only works on some desktop environments (e.g., GNOME-3 and maybe Ubuntu’s standard Unity interface). Also, according to https://www.x.org/wiki/RadeonFeature, examples of such AMD cards of the “Sea Islands” (CIK) generation are the models HD7790, R7 260, R9 290, and R9 M280X. These are marketing names that map to true hardware in often very confusing ways, so it is better to double-check. Such cards are also known under their internal GPU core code names BONAIRE, KABINI, MULLINS, KAVERI, HAWAII, or as cards of the “GraphicsCoreNext 1.1” (GCN 1.1) or “2nd generation graphics core next” generation. According to https://en.wikipedia.org/wiki/Graphics_Core, the Radeon HD 8770 may work as well. After updating to the latest Psychtoolbox and running PsychLinuxConfiguration, the system should be rebooted for the hdmi deep color support to enable. A “cat/sys/module/radeon/parameters/deep_color” must report “1” for deep color mode to be enabled. Next, verify Linux kernel’s extended debug output to make sure the projector is truly detected as 12 bpc capable, because even expensive display devices can have buggy EDID information which misrepresents their color resolution. One way to check this is to open a terminal and type “sudo su”, “echo 1 > /sys/module/drm/parameters/debug”, unplug/replug the projector to trigger the display redetection, “cat/var/log/syslog | grep bpc” (should report “… Display bpc=12, returned bpc=12” and “Assigning HDMI sink color depth as 12 bpc”), and “echo 0 > /sys/module/drm/parameters/debug)”.]

Kleiner M, Brainard D, Pelli D. What’s new in Psychtoolbox-3? Perception 36 ECVP Abstract Supplement; 2007.

Kothe C. Lab streaming layer (LSL); 2014 [accessed 2015 Oct 26]. https://github.com/sccn/labstreaminglayer.

Kremkow J, Jin J, Wang Y, Alonso JM. Principles underlying sensory map topography in primary visual cortex. Nature. 2016;533(7601):52.

Kummerer M, Wallis TS, Gatys LA, Bethge M. Understanding low- and high-level contributions to fixation prediction. Paper presented at the IEEE International Conference on Computer Vision; 2017; Venice, Italy.

Li Z. A neural model of contour integration in the primary visual cortex. Neural Comput. 1998;10(4):903–940.

Li Z. Neural circuit models for computations in early visual cortex. Current Opinion in Neurobiology. 2011;21(5):808–815.

Lim H, Wang Y, Xiao Y, Hu M, Felleman DJ. Organization of hue selectivity in macaque V2 thin stripes. Journal of Neurophysiology. 2009;102(5):2603–2615. doi:10.1152/jn.91255.2008.

Lipin-Dietz. Titmus i500 vision screener; 2012 [accessed 2019 July 12]. http://www.lipindietz.com/i500%20vision%20tester.htm.

Luck SJ. An introduction to the event-related potential technique. Cambridge (MA): MIT Press; 2014

Marathe AR, Files BT, Canady JD, Drnec KA, Lee H, Kwon H, Stump E. Heterogeneous systems for information-variable environments (HIVE). Aberdeen Proving Ground (MD): Army Research Laboratory (US); 2017. Report No.: ARL-TR-8027.

Moore PO, editor. Nondestructive testing handbook. Vol. 9 visual testing. 3rd ed. Columbus (OH): American Society for Nondestructive Testing, Inc.; 2010.

Motoyoshi I, Nishida SY, Sharan L, Adelson EH. Image statistics and the perception of surface qualities. Nature. 2007;447(7141):206.

Polat U, Sagi D. Lateral interactions between spatial channels: suppression and facilitation revealed by lateral masking experiments. Vision Research. 1993;33(7):993–999.

Polat U, Norcia AM. Neurophysiological evidence for contrast dependent long-range facilitation and suppression in the human visual cortex. Vision Research. 1996;36(14):2099–2109.

Polat U, Sagi D. Temporal asymmetry of collinear lateral interactions. Vision Research. 2006;46(6–7):953–960.

Polat U, Mizobe K, Pettet MW, Kasamatsu T, Norcia AM. Collinear stimuli regulate visual responses depending on cell’s contrast threshold. Nature. 1998;391(6667):580.

Radonjic A, Allred SR, Gilchrist AL, Brainard DH. The dynamic range of human lightness perception. Curr Biol. 2011;21(22):1931–1936. doi:10.1016/j.cub.2011.10.013

Ramsden BM, Hung CP, Roe AW. Real and illusory contour processing in area V1 of the primate: a cortical balancing act. Cereb Cortex. 2001;11(7):648–665.

Roe AW, Chelazzi L, Connor CE, Conway BR, Fujita I, Gallant JL, Vanduffel W. Toward a unified theory of visual area V4. Neuron. 2012;74(1):12–29.

Schroeder CE, Mehta AD, Givre SJ. A spatiotemporal profile of visual system activation revealed by current source density analysis in the awake macaque. Cereb Cortex. 1998;8(7):575–592.

Stewart CE, Hussey A, Davies N, Moseley MJ. Comparison of logMAR ETDRS chart and a new computerised staircased procedure for assessment of the visual acuity of children. Ophthalmic and Physiological Optics. 2006;26(6):597–601.

Tanigawa H, Wang Q, Fujita I. Organization of horizontal axons in the inferior temporal cortex and primary visual cortex of the macaque monkey. Cereb Cortex. 2005;15(12):1887–1899.

Töllner T, Zehetleitner M, Gramann K, Müller HJ. Stimulus saliency modulates pre-attentive processing speed in human visual cortex. PloS One. 2011;6(1):e16276.

Ts’o DY, Gilbert CD, Wiesel TN. Relationships between horizontal interactions and functional architecture in cat striate cortex as revealed by cross-correlation analysis. J Neurosci. 1986;6(4):1160–1170.

Ts’o DY, Roe AW, Gilbert CD. A hierarchy of the functional organization for color, form and disparity in primate visual area V2. Vision Research. 2001;41(10–11):1333–1349.

Ulrich R, Miller J. Using the jackknife-based scoring method for measuring LRP onset effects in factorial designs. Psychophysiology. 2001;38(5):816–827.

Wachtler T, Sejnowski TJ, Albright TD. Representation of color stimuli in awake macaque primary visual cortex. Neuron. 2003;37(4):681–691.

Wang Y, Xiao Y, Felleman DJ. V2 thin stripes contain spatially organized representations of achromatic luminance change. Cerebral Cortex. 2007;17(1):116–129. doi:10.1093/cercor/bhj131.

Zdravković S, Economou E, Gilchrist A. Grouping illumination frameworks. Journal of Experimental Psychology: Human Perception and Performance. 2012;38(3):776.

Zhou H, Friedman HS, Von Der Heydt R. Coding of border ownership in monkey visual cortex. Journal of Neuroscience. 2000;20(17):6594–6611.

Zucker SW, David C, Dobbins A, Iverson L. The organization of curve detection: Coarse tangent fields and fine spline coverings. Paper presented at the 2nd International Conference on Computer Vision; 1988; Tampa, FL.

